# Spatially Organized DNA-templated Silver Nanoclusters as Potent Antimicrobial Agents for ESKAPE Infections

**DOI:** 10.1101/2025.10.02.680010

**Authors:** Elizabeth Skelly, Krishna Majithia, Laura P. Rebolledo, Camila Fonseca Rizek, Silvia Figueiredo Costa, Alora R. Dunnavant, Cheyenne Vasquez, Alexander J. Lushnikov, Alexey V. Krasnoslobodtsev, Taejin Kim, Renata de Freitas Saito, Roger Chammas, M. Brittany Johnson, Kirill A. Afonin

## Abstract

Antibiotic-resistant bacteria cause more than one million deaths annually worldwide. The rapid evolution and gene exchange among pathogens often render new antibiotic drugs ineffective soon after deployment, underscoring the urgent need for alternative therapeutic strategies. Nanoscale silver is well known for its innate bacteriostatic and bactericidal activity but typically requires high concentrations for efficacy that causes toxicities and limits broader clinical applications. To address this, we introduce programmable self-assembling DNA scaffolds that template, stabilize, and spatially organize multiple copies of monodisperse silver nanoclusters (DNA-AgNCs). These assemblies enhance antimicrobial potency of formulations while also exhibiting intrinsic fluorescence, providing dual functionality for therapeutic and bioimaging applications. Detailed characterization identified DNA-AgNC scaffolds with improved stability and enhanced activity against clinically relevant antibiotic-resistant planktonic ESKAPE pathogens. We also revealed the DNA-AgNC constructs that significantly lowered intracellular infections of human bone cells with *Staphylococcus aureus*. Collectively, the results highlight spatially organized DNA-AgNCs as a promising modular platform for next-generation antibacterial therapy and real-time bioimaging.

---

The rise of multidrug-resistant bacterial species has made antibiotic resistance a pressing global health and economic crisis that must be addressed through the development of novel antibacterial therapies capable of halting the pathogenesis of systemic bacterial infections.^1, 2^ Among the most clinically significant organisms are ESKAPE pathogens, including *Enterococcus faecium* (*E. faecium*), *Staphylococcus aureus* (*S. aureus*), *Klebsiella pneumoniae* (*K. pneumoniae*), *Acinetobacter baumannii* (*A. baumannii*), *Pseudomonas aeruginosa* (*P. aeruginosa*), and *Enterobacter spp*., which are known for their ability to resist killing by currently available antibiotics. *S. aureus* is especially concerning due to its ability to cause a wide range of infections, from superficial skin and soft tissue infections to life-threatening diseases such as sepsis, endocarditis, and osteomyelitis.^3, 4^

Nearly 80% of osteomyelitis results from a severe manifestation of *S. aureus* infection and is associated with prolonged illness, disability, and in some cases, mortality.^5, 6^ Treatment is challenging due to the multiple strategies that *S. aureus* employs to evade immune defenses and antimicrobial therapies.^4, 7^ Externally, the bacteria form biofilms on necrotic bone tissue, creating a protective matrix and shielding itself from antibiotics and immune clearance.^8, 9^ Internally, *S. aureus* invades resident bone cells, including osteoblasts, osteocytes, and osteoclasts, as demonstrated by both in vitro and in vivo studies. Once internalized, bacteria can persist as an intracellular reservoir, evading killing by most host immune cells and antibiotics.^7, 10-13^ These features contribute to recurrent, and chronic osteomyelitis and highlight the urgent need for alternative therapeutic strategies targeting *S. aureus*.

One possible therapeutic strategy involves combining the therapeutic properties of silver, a historically renowned antibacterial agent, with nanotechnology, thus engineering novel nanoscale silver formulations for advancing modern disease prevention.^14, 15^ These silver nanoparticles possess antimicrobial modalities that bacteria have difficulty evading and are one of the most widely accepted antibacterial nanoagents both in a clinical setting and across a multitude of consumer products (medical implants, wound dressings, textiles, cosmetics, *etc*).^15-19^ Although novel and clinically-accepted, the applications of silver nanoparticles are limited by poorly understood mechanisms and potential toxicity to mammalian cells.^20-23^ Due to these concerns, the use of silver nanoparticles is currently restricted to surface-level or localized therapeutics only. They have not expanded as clinically acceptable treatment methods for bacterial infections such as osteomyelitis.

DNA-templated and -stabilized silver nanoclusters (DNA-AgNCs) are a new promising alternative therapeutic that could expand the applicability of nanoscale silver.^24-27^ These atomically precise hybrid biomolecule-metal nanostructures typically contain 5-30 silver atoms, giving them unique and tunable optical properties with advantageous antibacterial activity.^28-34^ The unique fluorescence of DNA-AgNCs arises from their ultrasmall size as well as the templating oligonucleotide’s structure and complimentary base interactions associated with the primary structure. The specific fluorescence color and excitation-emission patterns depend on the stabilizing single-stranded (ss)DNA oligomer sequence, size, shape, and the number of neutral silver atoms (Ag0) per cluster.^34-42^ The antibacterial properties of this therapeutic have been shown to be effective against both Gram-negative and Gram-positive bacteria.^21, 24^-^27, 43^ However, the application of DNA-AgNCs in treating bacterial infections in mammalian model systems remains unexplored, thus presenting a promising path for future research and discovery.

Our previously published study demonstrated that DNA-hairpins with varying cytosine counts in the hairpin loop displayed the strong antibacterial activity against *Escherichia coli*.^22, 24^ While DNA-AgNCs have been evaluated against various Gram-positive and Gram-negative bacteria, these studies have focused on planktonic bacteria and often lack data on specific resistant strains of effectiveness of DNA-AgNCs in infected mammalian cells.^25, 33, 44, 45^

In current study, we screened a panel of 11 distinct DNA-AgNCs, each incorporating varying numbers of single-stranded cytosines within hairpin (HP) structures ranging from C5 to C15, and identified the C13 HP as the most stable, optimal for bioimaging, and effective antimicrobial agent against clinically relevant, antibiotic-resistant bacterial strains. We demonstrated that combining multiple C13 HPs into single-stranded DNA scaffolds, as well as into multi-stranded fibrous DNA assemblies, significantly enhances the antimicrobial activity of formulations. Moreover, both C13 DNA-AgNCs and C13-based fibrous DNA-AgNCs were successfully delivered into mammalian cells, where they restricted intracellular bacterial burden. Collectively, these findings demonstrate that increasing the higher number of DNA-AgNCs within a single structure enhances antibacterial efficacy while their therapeutic concentrations can be optimized to minimize toxicity to mammalian cells.

## RESULTS AND DISCUSSION

### DNA-AgNCs display significant antimicrobial activity against planktonic S. aureus

DNA HPs with cytosine-rich loops has been demonstrated to template DNA-AgNCs, which have gained interest due to their fluorescent and antibacterial properties.^23, 46-49^ However, effective antibacterial activity typically requires high concentrations, which may also impact mammalian cells. To address this limitation, the goal of this work was to engineer DNA-AgNCs with increased antibacterial activity and reduced toxicity.

DNA HPs with C5-C15 in the loop were used as a scaffold to synthesize 11 distinct DNA-AgNCs (**Fig. 1a**). Each sample exhibited unique fluorescence under UV excitation (340 nm), following a reverse rainbow order: smallest HP loops (C5 and C6) produced faint blue fluorescence, while the largest HP loops (C13 to C15) produced red fluorescence (**Fig. 1b**). To evaluate their optical properties, 3D excitation-emission spectra were recorded 24 hours after synthesis, revealing that each sample possessed unique excitation and emission maxima as shown in (**Fig. 1c**). Across all samples, fluorescence was the most intense under yellow-to-red visible light excitation. To assess stability of the samples, the excitation-emission spectra were measured at two- and four-weeks post synthesis while storing the samples in the dark at 4°C. Over time, most emission peaks diminished with aging, while some designs showed better stability than others. For example, C13 maintained a consistent emission signal over time (**Fig. 1c**) indicating the most stable confirmation.

**Figure 1.**
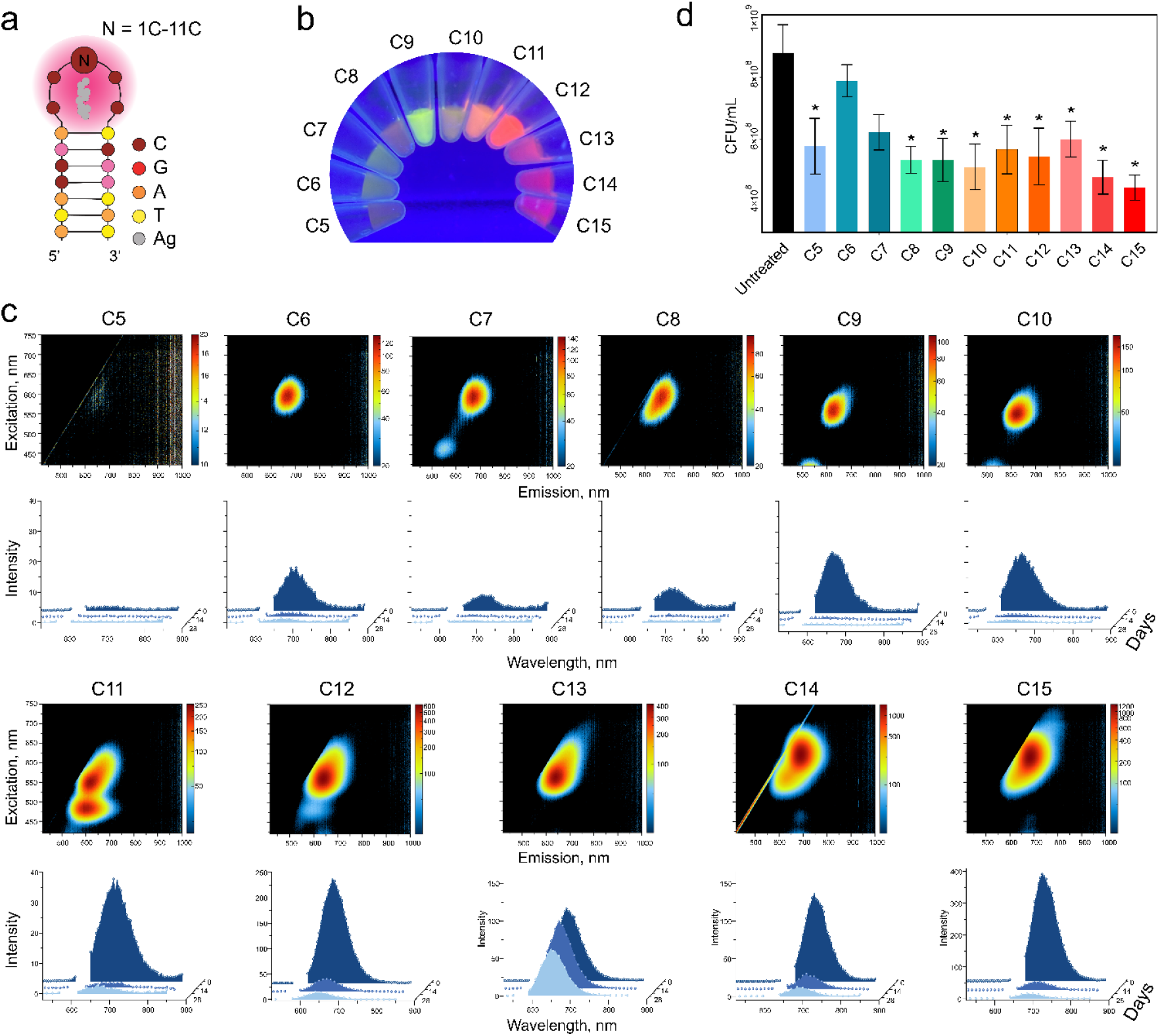
Evaluation of physical and biological property changes in DNA hairpins (HPs) with 5 to 15 cytosines in the loop (C5–C15). (**a**) Schematic representation of DNA HPs with a consistent stem region and varying numbers of cytosines. (**b**) Fluorescence intensity differences among DNA-AgNCs at 25 μM DNA. (**c**) Initial excitation-emission matrices and changes in fluorescence over four weeks for DNA-AgNCs at 10 μM DNA. (**d**) Colony-forming units (CFUs) of *S. aureus* treated with DNA-AgNCs at 4 μM DNA, assessed 6 hours post-treatment. Error bars represent mean ± SEM, n = 3, *P < 0.05.

Furthermore, the antibacterial activity of all DNA-AgNCs was compared against the clinically relevant *S. aureus* strain, UAMS-1. Among these, C5 and C8-C15 DNA-AgNC structures exhibited significant antibacterial effects compared to untreated controls (**Fig. 1d**), with no major differences observed between the structures. Interestingly, the degree of antibacterial activity did not show any correlation with a wide range of emissions. In contrast, C6 and C7 exhibited negligible antibacterial activity despite displaying higher fluorescence intensities in the excitation-emission spectra.

### Increasing the number of C13 hairpin loops on nanostructures enhances the antimicrobial activity against planktonic S. aureus

To enhance antibacterial activity and reduce the concentration required to kill *S. aureus*, we increased the number of DNA-AgNCs within a single nanostructure, thereby elevating the local concentration of silver. Based on the unique excitation emission spectra for each of the C5-15 hairpin structures and their relative stabilities, we selected the C13 hairpin to proceed with. The C13 hairpin has an emission in the red wavelength range which is ideal for bioimaging and maintains higher stability over time when compared to all other HPs.

We next tested multi-hairpin structures containing either two or three cytosine-rich loops, hereafter referred to as 2HP and 3HP, respectively. To assess the effect of structural flexibility, additional 3HP variants were designed with one, two, and three thymine spacers inserted between the adjacent HPs. Computational modeling supported these designs (**Fig. 2a-b**), while the single C13 hairpin was used as a reference control. Both 2HP and 3HP structures exhibited fluorescence comparable to that of the single C13 hairpin (**Fig. 2c-d**). Notably, the 3HP structure showed the highest stability among the constructs, with the smallest decrease in fluorescence over time.

**Figure 2.**
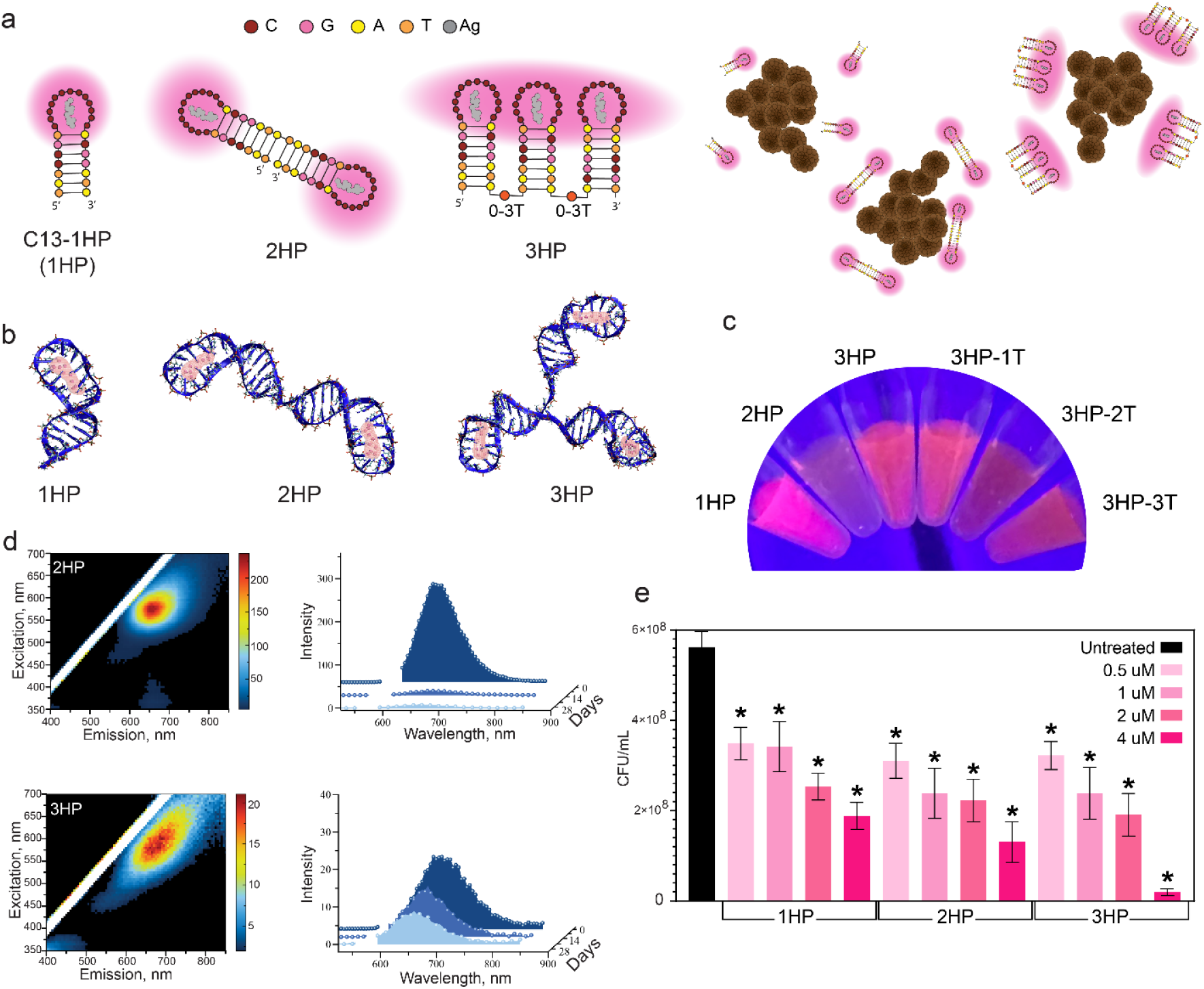
Evaluation of physical and biological property changes in multi-hairpin DNA-AgNCs. (**a**) Schematic of multi-hairpin DNA- AgNCs enabling more silver to surround bacteria at equimolar DNA concentration. (**b**) Computational models of multi-hairpin DNA- AgNCs. (**c**) Visual appearance of multi-hairpin DNA-AgNCs under UV excitation. (**d**) Initial excitation-emission matrices and fluorescence intensity changes over time of multi-hairpin DNA-AgNCs at 10 μM DNA. (**e**) Colony-forming units (CFUs) of *S. aureus* treated with multi-hairpin DNA-AgNCs at varying concentrations (0.5, 1, 2, 4 μM) of DNA, assessed 6 hours post-treatment. Error bars represent mean ± SEM, n = 9, *P < 0.05.

Next, we assessed if increasing the local concentration of silver influences antibacterial activity against planktonic *S. aureus* across a concentration range of 0.5 to 4 μM DNA. We observed a clear dose-dependent reduction in viable bacteria with maximal bacterial death at 4 μM DNA. Notably, samples containing a greater number of hairpin structures demonstrated a further reduction in bacterial viability at 4 μM DNA, indicating that higher localized silver loading enhances the antibacterial activity of these nanostructures (**Fig. 2e, Fig. S4**).

We further determined the minimal inhibitory concentration (MIC) and the minimal bactericidal concentration (MBC) of the 3HP DNA-AgNCs against clinically relevant antibiotic-resistant Gram-negative (*E. coli, P. aeruginosa and K. pneumoniae*) and Gram-positive (*S. aureus and E. faecalis*) bacterial species (**Fig. 3**, and **SI Fig. S3)**. All of the tested bacterial species are recognized as public health threats. According to the Center for Disease Control and Prevention (CDC) Report on Antibiotic Resistance Threats in the United States^50^, Carbapenem-resistant *Acinetobacter* is an urgent threat, while Vancomycin-resistant *E. faecalis* (VRE), Multidrug-resistant *P. aeruginosa* and Methicillin-resistant *S. aureus* (MRSA) are considered serious threats. Similarly, Carbapenem-resistant *K. pneumoniae* is included in the World Health Organization (WHO) list of emerging bacteria that pose the greatest threat to human health.^51^

**Figure 3.**
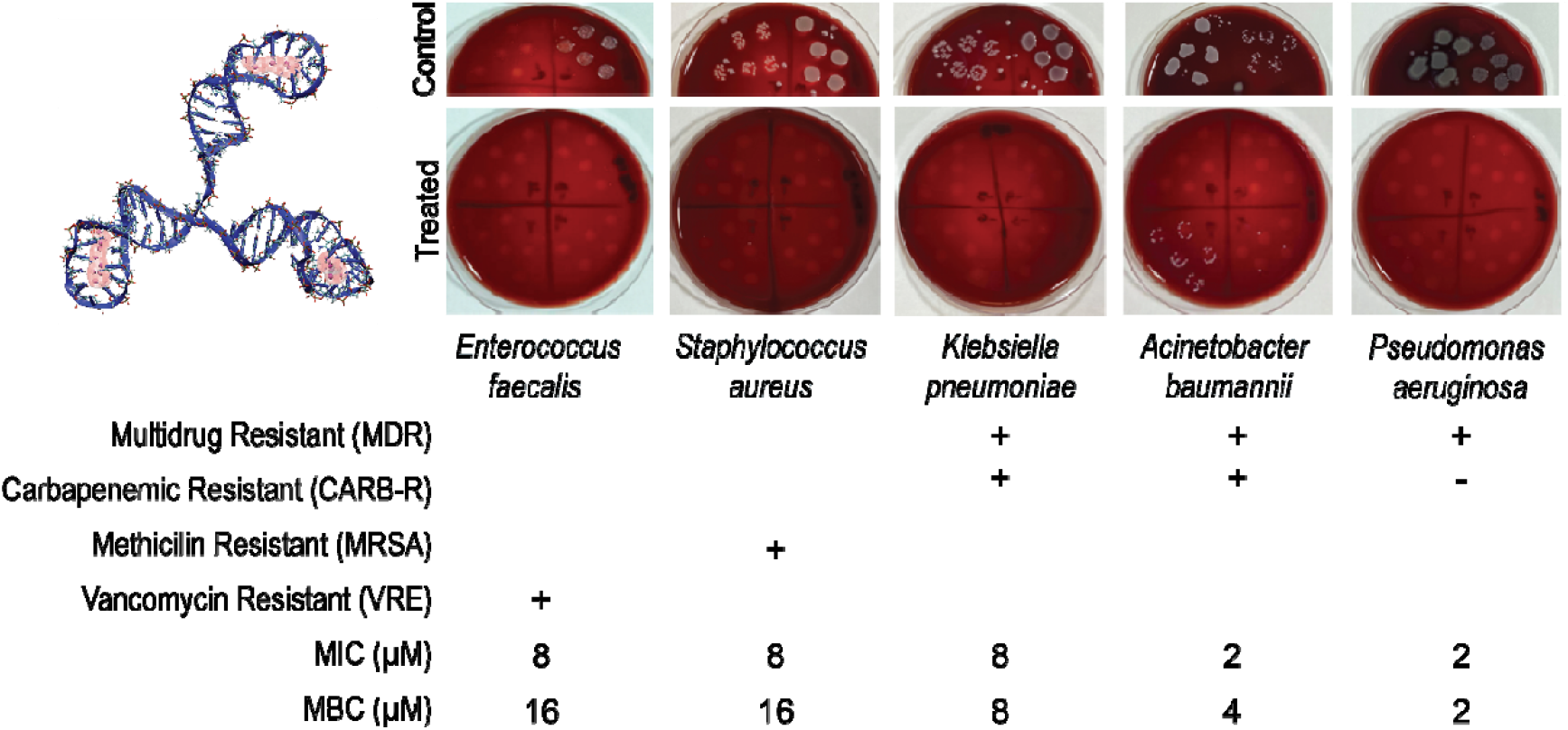
Evaluating C13-3HP DNA-AgNCs (3D model is shown in top left) against drug-resistant Gram-negative (*A. baumannii, P. aeruginosa, K. pneumoniae*) and Gram-positive (*S. aureus and E. faecalis*) bacteria, with representative plates for each control and treated bacteria used for counting viable CFU. The specific drug resistance is noted by (+), with minimum inhibitory concentrations (MIC) and minimum bactericidal concentrations (MBC) listed.

The 3HP DNA-AgNCs displayed broad antimicrobial activities against all antibiotic-resistant bacteria. Resistant clinical strains showed MICs of 2-4 μM for Gram-negative and 8-16 μM for Gram-positive isolates, except for *K. pneumoniae*, which exhibited a higher MIC of 8 μM compared to the Gram-negative ATCC strains. In contrast, susceptible ATCC strains displayed MICs of 1-4 μM for Gram-negative and 4-8 μM for Gram-positive bacteria (**SI Table S1**). Overall, Gram-positive bacteria showed higher MICs than Gram-negative. For MBCs, Gram-negatives matched their MICs across ATCC and clinical strains, whereas Gram-positives consistently required one dilution higher (8 μM for ATCC, 16 μM for resistant strains). This is consistent with previous studies demonstrating that silver nanoparticles were more effective against Gram-negative bacterial species.^52^ The observed differences in sensitivity of Gram-negative and Gram-positive bacteria may be in part due to the differences in bacterial structures. In contrast to Gram-negative bacteria, which have a thin peptidoglycan cell wall and an additional lipid outer membrane, Gram-positive bacteria have a thick peptidoglycan cell wall, which may provide a barrier hindering the permeation of Ag^+^ ions through the cytoplasmic membrane.^53, 54^ The consistent overlap of MIC and MBC values in Gram-negative bacteria suggests rapid bactericidal activity, while the one-dilution gap in Gram-positives implies a slower or less efficient killing process, possibly due to cell wall barriers limiting the effective intracellular accumulation of the nanoclusters.

### Fibrous DNA-AgNCs display enhanced antimicrobial activity against planktonic S. aureus

After the successful synthesis and characterization of multiple hairpin DNA-AgNCs, we engineered fibrous DNA-AgNCs. Fibrous DNA-AgNCs were made with either 1 or 2 hairpins per monomer, each mimicking the C13 single hairpin structure (**Fig. 4a, d**). Additionally, the fibrous DNA- AgNCs were designed to have varying degrees of flexibility with thymine linkers inserted between the adjacent HPs monomers, and the resulting structures were computationally modeled (**Fig. 4a, d**). Similar to the single and multi-hairpin C13 constructs, the fibrous DNA- AgNCs exhibited red fluorescence (**Fig. 4b**) maintaining their potential use for bioimaging applications. Notably, our data support an inverse relationship between structure flexibility and fluorescent stability. Stability testing showed that fibrous DNA-AgNCs with a single hairpin on each monomer had the most consistent excitation-emission spectra over time. The fibrous DNA- AgNCs with two hairpins on each monomer with the least amount of flexibility were found to be the most stable overtime as indicated by the excitation-emission spectra (**Fig. 4c**). AFM imaging confirmed the formation of fibrous DNA-AgNCs (**Fig. 4e**), consistent with the predicted designs.

**Figure 4.**
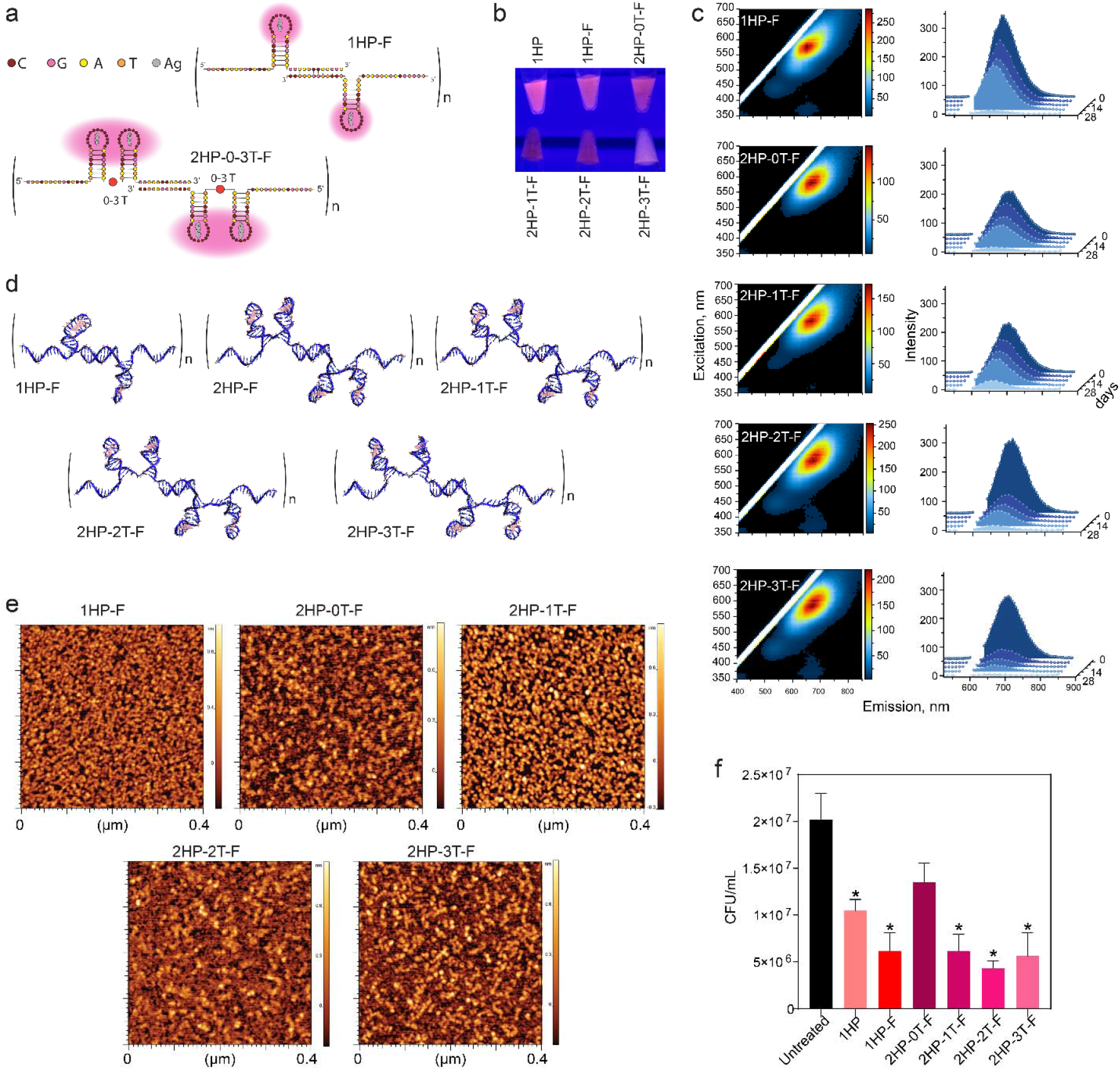
Evaluating the change in physical properties and biological activity of DNA-AgNCs when hairpin structures are introduced on fiber structures. (**a**) The depiction of fibrous DNA-AgNCs that either have a single hairpin on each monomer (e.g., 1HP-F) or two hairpins on each monomer with 0-3 thymines between the two fibers (e.g., 2HP-0T-F). (**b**) The colors of fibrous DNA-AgNCs when excited with UV light, as compared to 1HP DNA-AgNC. (**c**) Excitation-emission matrices for fibrous DNA-AgNCs at 10 μM DNA. (**d**) Computational models of fibrous DNA-AgNCs. (**e**) AFM images of the fibrous DNA-AgNCs. (**f**) Colony-forming units (CFUs) of *S. aureus* when treated with fibrous DNA-AgNCs at 0.5 μM DNA, analyzed 6 hours post-treatment. Error bars represent mean ± SEM, n = 3, *P < 0.05.

Following physical characterization, the antibacterial properties of fibrous DNA-AgNCs were tested against planktonic *S. aureus* (**Fig. 4f**) demonstrating stronger bactericidal activity compared to HP DNA-AgNCs. No significant differences in activity were observed among the various fibrous DNA-AgNCs. Thus, incorporating multiple hairpins on a single monomer was not advantageous over a single hairpin per monomer. Similarly, altering flexibility did not further enhance antibacterial efficacy.

### Fibrous DNA-AgNCs restrict intracellular S. aureus in resident bone cells

*S. aureus* is known to survive in both the extracellular and intracellular environments during osteomyelitis. It is now documented that resident bone cells, including bone-forming osteoblasts, serve as reservoirs of viable *S. aureus*. In the absence of a delivery reagent DNA-AgNCs remained extracellular. To deliver DNA-AgNCs to primary murine osteoblasts, nanostructures were complexed with Lipofectamine 2000 (L2K). The 1HP DNA-AgNCs and fibrous DNA-AgNCs were successfully delivered intracellularly to osteoblasts and visualized via fluorescent microscopy (**Fig. 5, S6**). Importantly, delivery of all DNA-AgNCs to *S. aureus*-infected osteoblasts significantly reduced intracellular bacterial burden. Our results indicate that all tested DNA-AgNCs reduced *S. aureus* intracellular CFU counts compared to untreated and carrier-only controls, with the 2HP-2T and 2HP-3T fibrous DNA-AgNCs demonstrating the greatest efficacy. Although these constructs did induce an inflammatory response in osteoblasts in IL-6, we did not observe any off-target toxicity and found that immunostimulation is independent of TLR-9 activation of the NF- κB pathway (**Fig. S5**). Importantly, at the concentration tested in primary murine osteoblasts we were able to significantly reduce bacterial burden in the absence of any undesired mammalian cell death.

**Figure 5.**
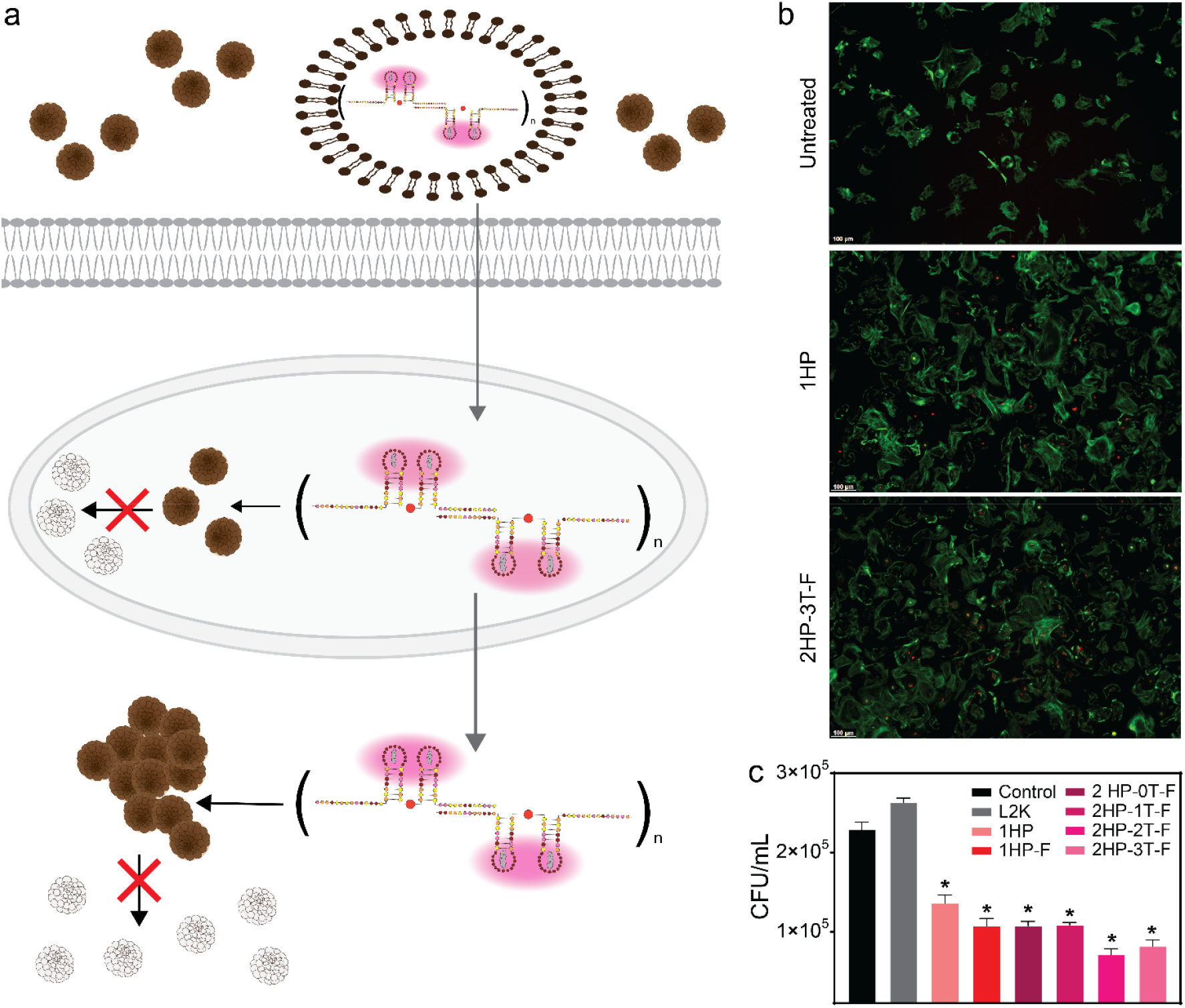
a. Primary murine osteoblasts were first infected with *S. aureus* (MOI 75:1), then transfected with DNA-AgNCs (at 0.5 μM DNA) using Lipofectamine 2000 (L2K). (**b**) Uptake images of osteoblasts at 2 hours post-transfection with DNA-AgNCs. More images can be found in **Fig. S6**. (**c**) CFU of *S. aureus* at 6 hours-post infection for various DNA-AgNCs. Error bars represent mean ± SEM, n = 3, *P < 0.05.

In conclusion, DNA-AgNCs hold significant promise as antibacterial therapeutics with bioimaging capabilities. Since DNA sequence and structure can be precisely tuned, DNA- AgNCs can be engineered for specific purposes, thereby optimizing stability, fluorescence properties, and antimicrobial activity. Our work demonstrates that C13-3HP DNA-AgNCs are effective antimicrobials against antibiotic-resistant pathogens. Specifically, we show that the C13 hairpin has enhanced stability while possessing optimal fluorescence for bioimaging. Importantly, the C13 hairpin exhibits antimicrobial activity against clinically relevant Gram-negative and Gram-positive bacterial strains that display antibiotic resistance. The fibrous DNA- AgNCs containing multiple C13 hairpins increase the delivery concentration of silver ions, further enhancing antimicrobial activity. Importantly, C13 hairpins and fibrous DNA-AgNCs can be delivered in the absence of a carrier to target extracellular bacterial burden or complexed with a lipid barrier to significantly restrict intracellular burden. However, intracellular use will require careful dose optimization to minimize cytotoxicity and possible immune stimulation. Collectively, these results establish the therapeutic potential of highly modular bi-functional DNA-AgNC platform as alternatives to antibiotics, recognizing the need for further experimental studies to investigate the mechanisms underlying the antimicrobial activity of DNA-AgNCs and assessing their efficiency in vivo.

## Materials and Methods

### DNA-AgNCs C5-15 Hairpin Synthesis

To create each unique DNA-AgNC, AgNO_3_ was mixed with the specified DNA template, which yielded a final DNA concentration of 25 μM, in 4 Mm NH_4_OAc, with a pH of 6.9, providing a 1:1 ratio of Ag^+^:Cytosine. Each sample solution was vortexed, centrifuged, and heated to 95°C for 2 minutes. Then, samples were immediately incubated in an ice bath for 20 minutes. During each incubation period, a fresh 10 mM NaBH_4_ solution was made for the individual DNA-AgNC syntheses using cold water. To reduce Ag^+^, after incubation, an equimolar amount of NaBH_4_ was added to each sample, mixed by pipetting and quick centrifugation, and allowed to sit at 4 °C for a minimum of 15 hours. Then, excess reagents were removed by washing the newly formed DNA-AgNCs in a 3 kDa molecular weight cut-off centrifuge filter with 20% initial volume of 4 mM NH_4_OAc, spun 3 times at 12,000 RCF for 25 minutes at 4°C. Estimated concentrations of the pure DNA-AgNC solutions were quantified by measuring the absorption of 260 nm light using Nanodrop 2000 and diluted to the concentration of interest. All DNA sequences are reported in Supporting Information, **Table S1**.

### Fibrous DNA-AgNCs

Equimolar amounts of complementary fiber strands A and B were incubated separately at 95°C for 5 minutes. After strand incubation, the complementary strands were added to AgNO_3_ and 20 mM NH_4_Oac (final concentration of 4 mM NH_4_OAc), then incubated at 25°C for 20 minutes. During each incubation period, a fresh 10 mM NaBH_4_ solution, using cold double-deionized water, was made. To reduce Ag^+^, after incubation, an equimolar amount of NaBH_4_ was added to each sample, mixed by pipetting, and allowed to sit at 4°C for a minimum of 15 hours. Fiber samples were washed following the hairpin washing protocol. The melting temperature (36°C) of the strands and 4°C was also evaluated to find the best temperature for synthesis. 25°C was determined to be the best temperature for the second incubation because the DNA-AgNC run on 8% native polyacrylamide gel electrophoresis was the most similar to that of the DNA fiber (**Fig. S1**).

### Building atomic models of C13-Ag_10_ HP structures

The stem region of DNA was built by Accelrys Discovery Studio. An initial hairpin (HP) loop structure was built based on an NMR DNA HP loop (PDB ID: 1JVE).^55^ The first base in the HP loop was replaced by cytosine using Accelrys Discovery Studio. From the second to the last base was replaced by twelve cytosine bases associated with Ag_10_ cluster, whose atomic conformation was reported in our previous study.^56^ Energy minimization was applied to the assembled C13-Ag_10_ structures to refine the atomic geometry using CHARMM36 nucleic acids force fields and Ag atom force fields.^57^ Energy energy-minimized C13-Ag_10_ structure was used to build C13-Ag_10_ structures and C13- Ag_10_ fibers. The energy minimizations were applied to C13-Ag_10_ structures and C13-Ag_10_ fibers.

### Fluorescence Measurements

Using a 96-well plate, 100 μL of each sample at a concentration of 10 μM was pipetted into each well. The plate was loaded into a Tecan Spark microplate reader, and 3D excitation-emission matrices were recorded. The excitation data were measured over a range of 350-700 nm with a manual gain of 150 and a 5 nm step size between measurements. The emission data were also measured with a manual gain of 150, but over a range of 400-850 nm with a 5 nm step size between measurements. Once all the data were recorded, it was plotted using MagicPlot Pro. These measurements were taken weekly or bi- weekly for 4 weeks to evaluate the change in fluorescence over time.

### Determination of minimum inhibitory concentration (MIC)

3HP DNA-AgNCs were tested against American Type Culture Collection (ATCC) standard strains (*E. coli* 25922, *P. aeruginosa* 27853, *K. pneumoniae* 13883, *S. aureus* 29213, and *E. faecalis* 29212). The resistant bacterial isolates belonged to the Bacteriology Laboratory LIM49 strains collection and were isolated from bloodstream infections of Hospital das Clínicas, Faculdade de Medicina, Universidade de São Paulo, São Paulo. The multi-resistant strains tested were *A. baumannii, P. aeruginosa, K. pneuminae, S. aureus* (MRSA), and *E. faecalis* (VRE). Each strain was cultivated aerobically on blood agar plates at 37°C for 24h to perform the susceptibility tests. MIC was determined by broth microdilution test using Mueller–Hinton Broth -MHB II (Difco™, BD, USA) for the 3HP DNA-AgNCs according to Clinical and Laboratory Standards Institute (CLSI) for antimicrobial susceptibility testing.^58^ The MIC of 3HP DNA-AgNCs was determined against five ATCC reference strains (EC 25922, PA 27853, KP 13883, SA 29213, and EF 29212) in duplicates and in three independent experiments. The 96-well assay plates containing different concentrations of 3HP DNA-AgNCs ranged from 8 to 0.0025 μM and were obtained by two-fold serial dilution and incubated at 35 ± 2 °C for 16 to 18 h, and the presence of turbidity on each concentration was visually determined with observation under transmitted light. Negative control was inoculated under identical conditions but without the addition of 3HP DNA-AgNCs, and sterility control was performed with MHB II without bacterial inoculation. The results were read by evaluating the turbidity of each well visually. The first well with no turbidity was defined as the MIC, expressed in μM. We also performed the same experiment with only the 3HP DNA-AgNCs diluent and determined that it did not interfere with the MICs, as all bacteria grew in all the diluent concentrations tested. (**Fig. S2, S3**).

### Determination of minimum bactericidal concentration (MBC)

To determine the MBC, 10μl of each well showing no obvious bacterial growth on the MIC test was seeded onto blood agar plates by streaking and incubated at 37°C for 24 hours. Subsequently, plates were examined, and the lowest concentration at which no visible growth was observed was taken as the MBC of 3HP DNA-AgNCs.

### Mammalian cell viability assays

To evaluate cell viability after transfection, an MTS assay was conducted using a 96-well flat-bottom Greiner plate. A total of 20 μL of MTS reagent was added to 100 μL of cells per well. The absorbance was measured at a wavelength of 638 nm using a Tecan Spark plate reader. Each condition was tested in biological triplicates, and the results were averaged and normalized to the cells-only control for analysis.

### S. aureus propagation

*S. aureus* strain UAMS-1 was grown on Luria broth (LB) agar plates overnight, followed by incubation in LB broth at 37ºC and 5% CO2 overnight. The number of colony-forming units (CFUs) was determined using a Genespec3 spectrophotometer as previously described (MiraiBio Inc.).

### Bacterial viability

*S. aureus* was seeded at a density of 1 × 10^6^ CFUs per well in a 96-well plate. Cells were left untreated or treated with the indicated concentrations of DNA-AgNCs in LB broth for 6 hours on an orbital shaker at 37ºC and 5% CO_2_. At 6 hours post-incubation, serial dilutions were performed, and CFUs were plated on LB agar plates overnight. The number of viable colonies was assessed by colony counting.

### Evaluating mammalian cell toxicity and immunostimulation

HEK-Blue hTLR9 cells were cultured according to InvivoGen’s protocols at 37 °C and 5% CO_2_ in DMEM supplemented with 10% heat-inactivated FBS, 1% penicillin–streptomycin, 100 μg/mL Normocin, 100 μg/mL Zeocin, and 10 μg/mL Blasticidin. For reverse transfection in 96-well plates, ∼80,000 cells per well were seeded onto preloaded treatments or controls. Poly I:C (5 μg/mL) was used as a positive control for SEAP activation in the Quanti-Blue assay. Intracellular delivery of DNA- AgNC treatments (4 μM) was achieved by pre-complexing DNA-AgNCs with L2K at room temperature for 30 min before transfection. After 24 h, SEAP activation and cell viability were assessed following the manufacturer’s protocols. For SEAP detection, 20 μL of cell supernatant was incubated for 2 h with 180 μL Quanti-Blue solution, and absorbance was measured at 638 nm using a Tecan Spark plate reader. Cell viability was evaluated using an MTS colorimetric assay, performed by adding a reagent equivalent to 20% of the final well volume, followed b ya 2 h incubation, with absorbance recorded at 490 nm. All assays were performed in biological triplicate. Results were averaged, normalized to untreated (cell-only) controls, and expressed as fold induction. Significance was calculated in GraphPad Prism using a two-way ANOVA.

### Isolation and maintenance of primary bone cells

Whole calvaria were harvested from 2-3-day-old neonatal mice and differentiated for 10 days as previously described.^59-63^ Differentiation was validated by alkaline phosphatase staining using a commercial kit (Abcam) and confirmed by microscopy, following established protocols.^63^

### Immunofluorescence microscopy of transfected osteoblasts

Osteoblasts were transfected with DNA-AgNCs using Lipofectamine 2000. Cells were fixed at 2h post-transfection and processed for immunofluorescence microscopy for actin using Alexa Fluor 488 Phalloidin (ThermoFisher Scientific, A12379, 100 nM). Cells were imaged using a Leica DMi8 fluorescence microscope.

### Bacterial infection of osteoblasts

Primary murine osteoblasts were seeded at a density of 1 × 10^6^ cells per well and infected with *S. aureus* at a multiplicity of infection (MOI) of 75 bacteria per host cell in antibiotic-free media for 2 hours. Following infection, the medium was replaced with fresh medium containing 1% penicillin-streptomycin. At 8 hours post-infection, cell supernatants were collected for analysis. Additionally, osteoblasts were lysed using Saponin, and intracellular CFUs were plated on LB agar plates overnight. The number of viable colonies was assessed by colony counting.

### DNA-AgNCs transfection of osteoblasts

Primary murine osteoblasts were seeded at a density of 1 × 10^6^ cells per well and infected with *S. aureus* as previously described. Following infection, cells were transfected with 0.5 μM DNA-AgNCs using the lipid-based carrier, L2K (Invitrogen), in antibiotic-free medium for 2 hours. Following transfection, fresh medium containing 1% penicillin-streptomycin was added. At 8 hours post-infection, cell supernatants were collected for analysis. Additionally, osteoblasts were lysed using Saponin, and intracellular CFUs were plated on LB agar plates overnight. The number of viable colonies was assessed by colony counting.

### Enzyme-linked immunosorbent assays

Specific capture enzyme-linked immunosorbent assays (ELISAs) were performed to measure osteoblast production of interleukin-6 (IL-6) in response to *S. aureus* infection and transfection of nanoparticles. IL-6 production was determined using commercially available antibody pairs (BD Biosciences; 554400, 554402). Recombinant proteins for each target were used to generate standard curves, and protein concentrations were determined by matching sample absorbance values to the corresponding standard curve.

## ACKNOWLEDGEMENTS

The research reported in this publication was supported by the National Institute of General Medical Sciences of the National Institutes of Health under Award Number R35 GM139587 (to K.A.A.). The project was also supported in part by R21 AI190691-01 (to M.B.J and K.A.A.). The content of this publication does not necessarily reflect the views or policies of the Department of Health and Human Services, nor does mention of trade names, commercial products, or organizations imply endorsement by the US Government. This material is based upon work supported in part by the North Carolina Biotechnology Center (2025-FLG-0142). We thank Dr. Lewis A. Rolband for designing the original 2HP and 3HP oligonucleotides.

## TOC

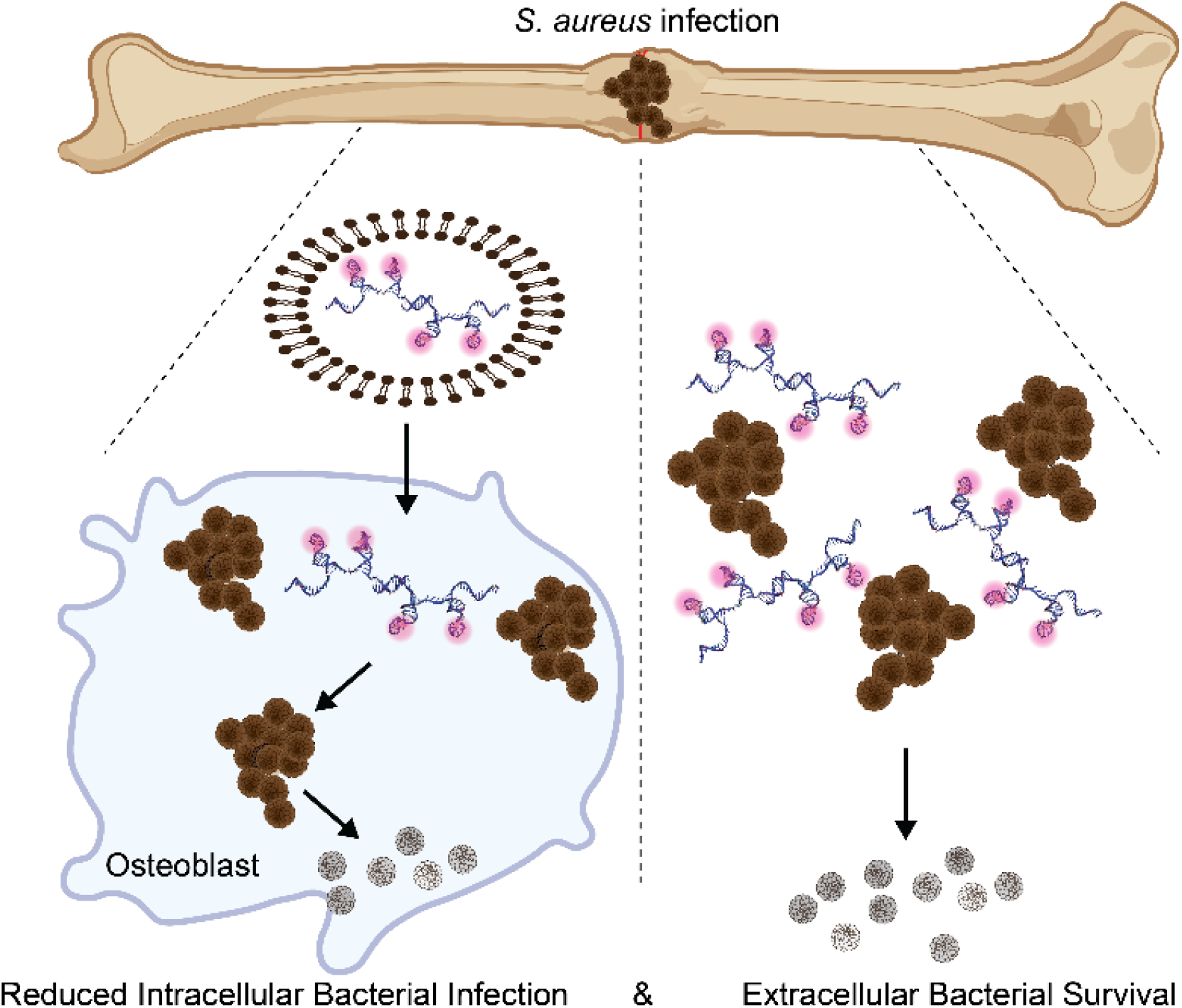

