## Supporting Information, Table S1, Fig. S1, Fig. S2, S3, Fig. S4, Fig. S5, Fig. S6, for "Spatially Organized DNA-templated Silver Nanoclusters as Potent Antimicrobial Agents for ESKAPE Infections"

**Sequences used in this project**

| **Complex Type** | **Strand Name** | **Sequence (5’—3’)** |
| --- | --- | --- |
| Single HP | C5 | TATCCGTCCCCCACGGATA |
| Single HP | C6 | TATCCGTCCCCCCACGGATA |
| Single HP | C7 | TATCCGTCCCCCCCACGGATA |
| Single HP | C8 | TATCCGTCCCCCCCCACGGATA |
| Single HP | C9 | TATCCGTCCCCCCCCCACGGATA |
| Single HP | C10 | TATCCGTCCCCCCCCCCACGGATA |
| Single HP | C11 | TATCCGTCCCCCCCCCCCACGGATA |
| Single HP | C12 | TATCCGTCCCCCCCCCCCCACGGATA |
| Single HP | C13 (1HP) | TATCCGTCCCCCCCCCCCCCACGGATA |
| Single HP | C14 | TATCCGTCCCCCCCCCCCCCCACGGATA |
| Single HP | C15 | TATCCGTCCCCCCCCCCCCCCCACGGATA |
| Multiple HP | 2HP | TATCCGTCCCCCCCCCCCCCACGGATATATCCGTCCCCCCCCCCCCCACGGATA |
| Multiple HP | 3HP | TATCCGTCCCCCCCCCCCCCACGGATATATCCGTCCCCCCCCCCCCCACGGATAACGGATACCCCCCCCCCCCCTATCCGT |
| Multiple HP | 3HP-1T | TATCCGTCCCCCCCCCCCCCACGGATATTATCCGTCCCCCCCCCCCCCACGGATATACGGATACCCCCCCCCCCCCTATCCGT |
| Multiple HP | 3HP-2T | TATCCGTCCCCCCCCCCCCCACGGATATTTATCCGTCCCCCCCCCCCCCACGGATATTACGGATACCCCCCCCCCCCCTATCCGT |
| Multiple HP | 3HP-3T | TATCCGTCCCCCCCCCCCCCACGGATATTTTATCCGTCCCCCCCCCCCCCACGGATATTTACGGATACCCCCCCCCCCCCTATCCGT |
| Fiber | 1HP-F-A | GTTCATCTGCACCAACGGATACCCCCCCCCCCCCTATCCGTGGAATCCAAGGA |
| Fiber | 1HP-F-B | TGGTGCAGATGAACACGGATACCCCCCCCCCCCCTATCCGTTCCTTGGATTCC |
| Fiber | 2HP-0T-F-A | GTTCATCTGCACCAACGGATACCCCCCCCCCCCCTATCCGTTATCCGTCCCCCCCCCCCCCACGGATAGGAATCCAAGGA |
| Fiber | 2HP-0T-F-B | TGGTGCAGATGAACACGGATACCCCCCCCCCCCCTATCCGTTATCCGTCCCCCCCCCCCCCACGGATATCCTTGGATTCC |
| Fiber | 2HP-1T-F-A | GTTCATCTGCACCAACGGATACCCCCCCCCCCCCTATCCGTTTATCCGTCCCCCCCCCCCCCACGGATAGGAATCCAAGGA |
| Fiber | 2HP-1T-F-B | TGGTGCAGATGAACACGGATACCCCCCCCCCCCCTATCCGTTTATCCGTCCCCCCCCCCCCCACGGATATCCTTGGATTCC |
| Fiber | 2HP-2T-F-A | GTTCATCTGCACCAACGGATACCCCCCCCCCCCCTATCCGTTTTATCCGTCCCCCCCCCCCCCACGGATAGGAATCCAAGGA |
| Fiber | 2HP-2T-F-B | TGGTGCAGATGAACACGGATACCCCCCCCCCCCCTATCCGTTTTATCCGTCCCCCCCCCCCCCACGGATATCCTTGGATTCC |
| Fiber | 2HP-3T-F-A | GTTCATCTGCACCAACGGATACCCCCCCCCCCCCTATCCGTTTTTATCCGTCCCCCCCCCCCCCACGGATAGGAATCCAAGGA |
| Fiber | 2HP-3T-F-B | TGGTGCAGATGAACACGGATACCCCCCCCCCCCCTATCCGTTTTTATCCGTCCCCCCCCCCCCCACGGATATCCTTGGATTCC |

Cytosine-rich hairpin sections are underlined.

**Supporting Figures**

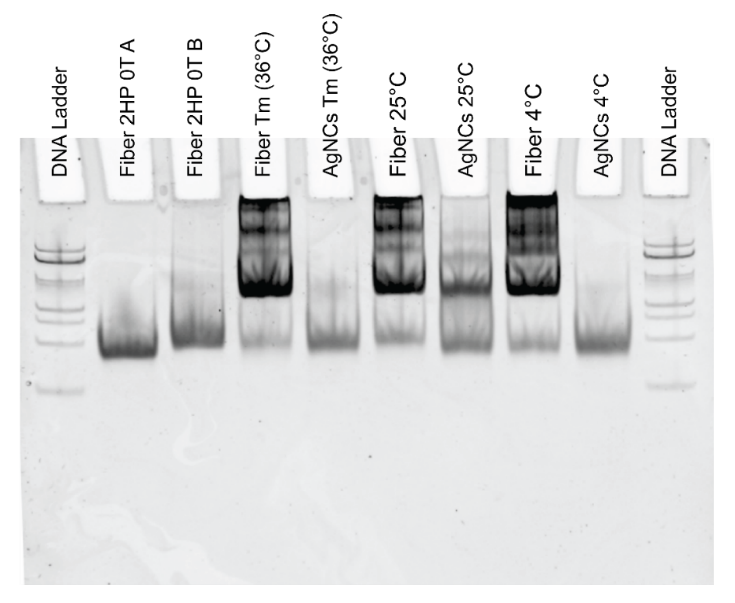

**Figure S1.** Evaluating Fiber DNA-AgNCs assembly incubation temperatures. Fiber DNA-AgNCs were found to have the most similar structure to Fiber DNA when incubated at 25°C, whereas the structures incubated at 36°C and 4°C look more similar to individual monomers. Thus, the Fiber DNA-AgNCs were synthesized with the incubation step at 25°C.

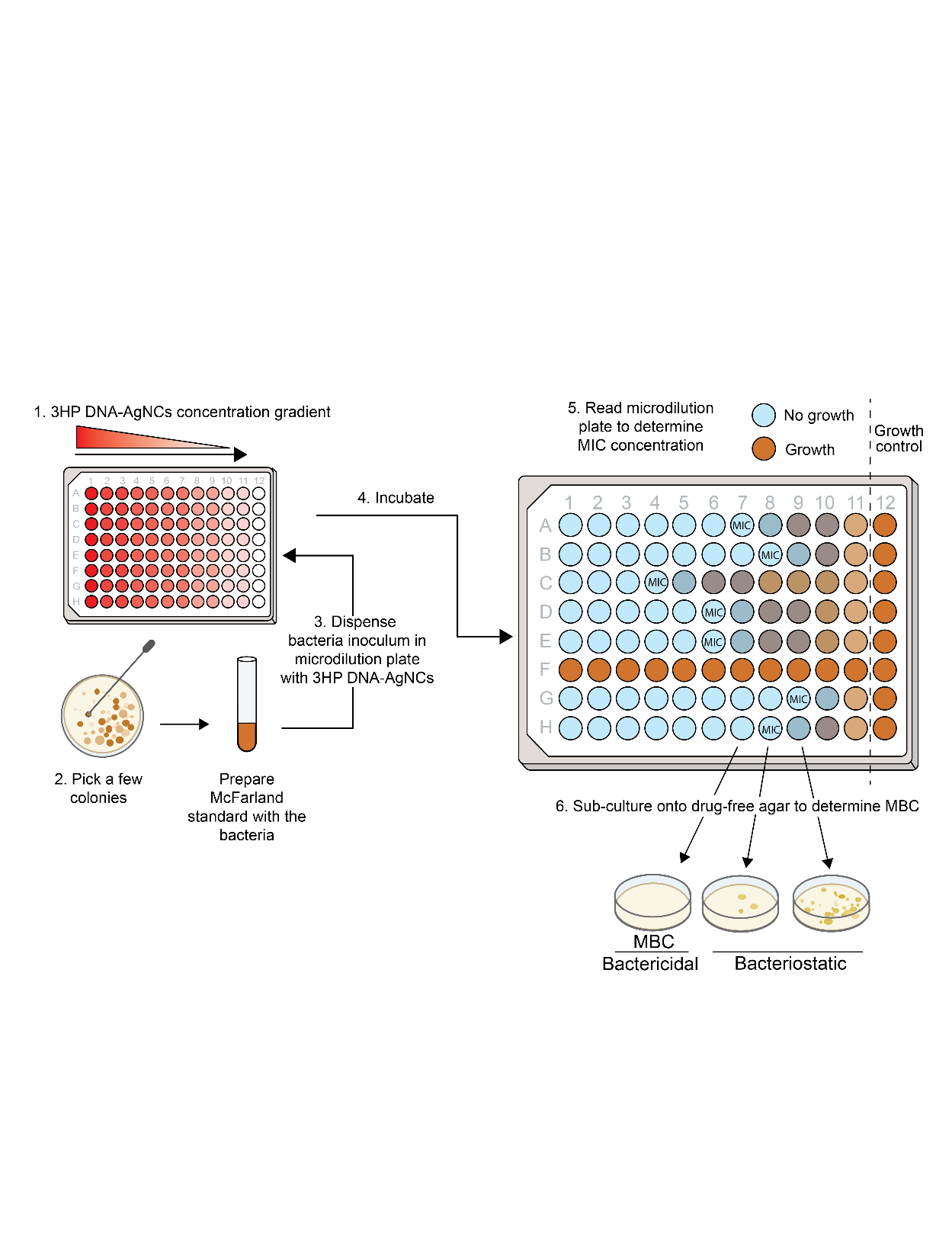

**Figure S2.** Scheme of MIC/MBC assay performed according to CLSI (Clinical and Laboratory Standards Institute) guidelines, (1) first prepare a serial dilution of C13-3HP DNA-AgNCs in a microdilution plate. (2) Prepare the inoculum by taking a few colonies from an agar plate with a sterile swab, prepare a 0.5 McFarland standard, and dilute the McFarland standard into the media. (3) Dispense the inoculum into the microdilution plate with the serial diluted C13-3HP DNA-AgNCs and incubate the plate at 37°C. (4) Read the plate to determine the MIC value. (5) Plate a portion of each well on an agar media, incubate and check for colonies to determine the MBC.

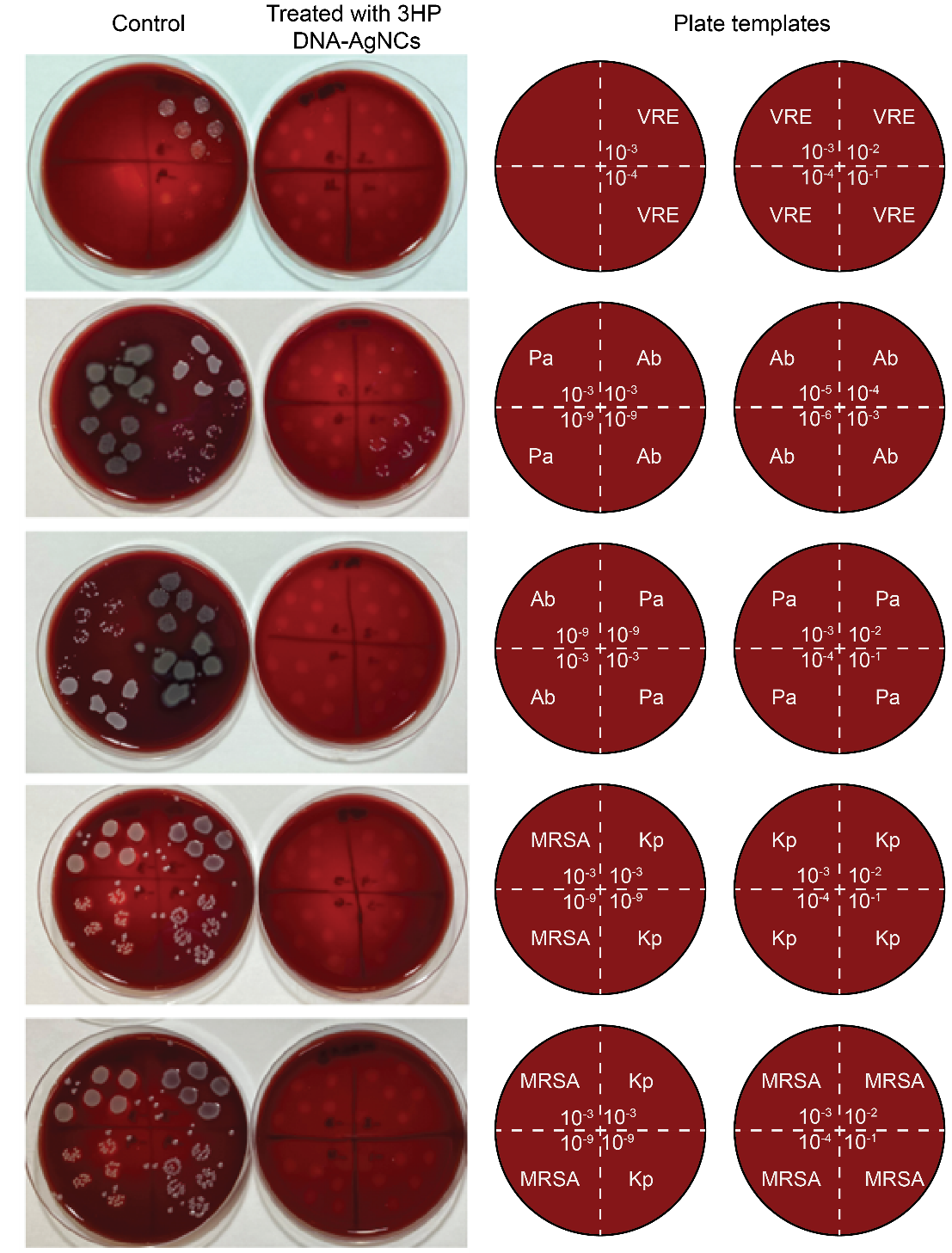

**Figure S3.** Measure the antimicrobial effects of 3HP DNA-AgNCs on antibiotic-resistant strains. After determining MIC, an aliquot was removed and up to 9 serial ten-fold dilutions were performed in Mueller–Hinton Broth to obtain 10^-1^ through 10^-9^ dilutions. An aliquot of 10 µl of each dilution was spotted five times onto a blood agar plate divided into four quadrants. After 24 hours of incubation at 37°C, colonies were observed. In the dilutions of control without 3HPs DNA-AgNCs, it was not possible to count single colonies, precluding the determination of CFU/mL. Legend: VRE (Vancomycin-resistant Enterococcus faecalis), Kp (Multidrug-resistant and Carbapenemic-resistant *Klebsiella pneumoniae*), Ab (Multidrug-resistant and Carbapenemic-resistant *Acinetobacter baumannii*), Pa (Multidrug-resistant *Pseudomonas aeruginosa*), and MRSA (Methicillin-resistant *Staphylococcus aureus*).

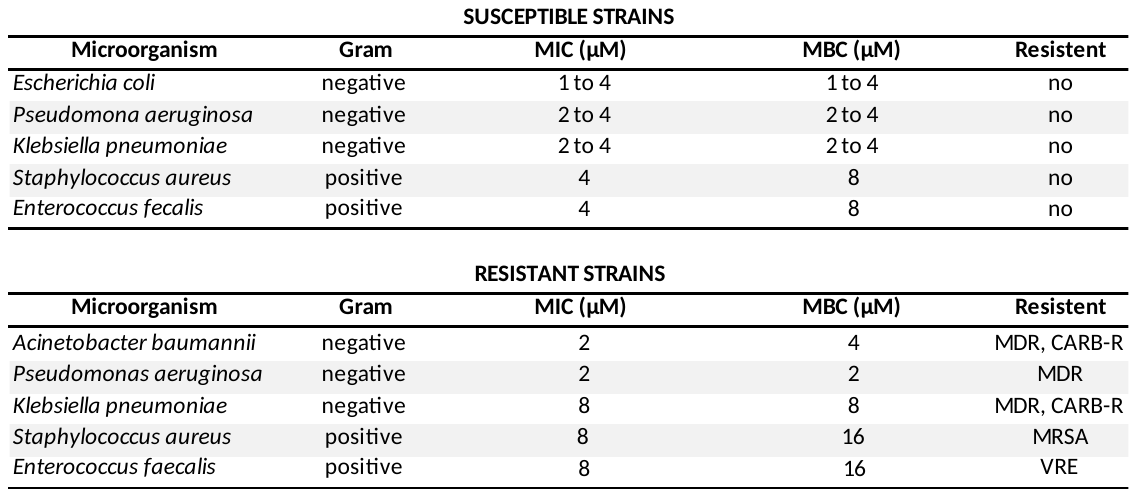

**Table S1.** Minimum inhibitory concentration/minimum bactericidal concentration *E. coli* 25922, *P. aeruginosa* 27853, *K. pneumoniae* 13883, *S. aureus* 29213, and *E. faecalis* 29212. Minimum inhibitory concentration/minimum bactericidal concentration for resistant bacterial isolates of Gram-negative (*Acinetobacter baumannii*, *Pseudomonas aeruginosa*, and *Klebsiella pneumoniae*) and Gram-positive (*Staphylococcus aureus* and *Enterococcus faecalis*) bacteria.

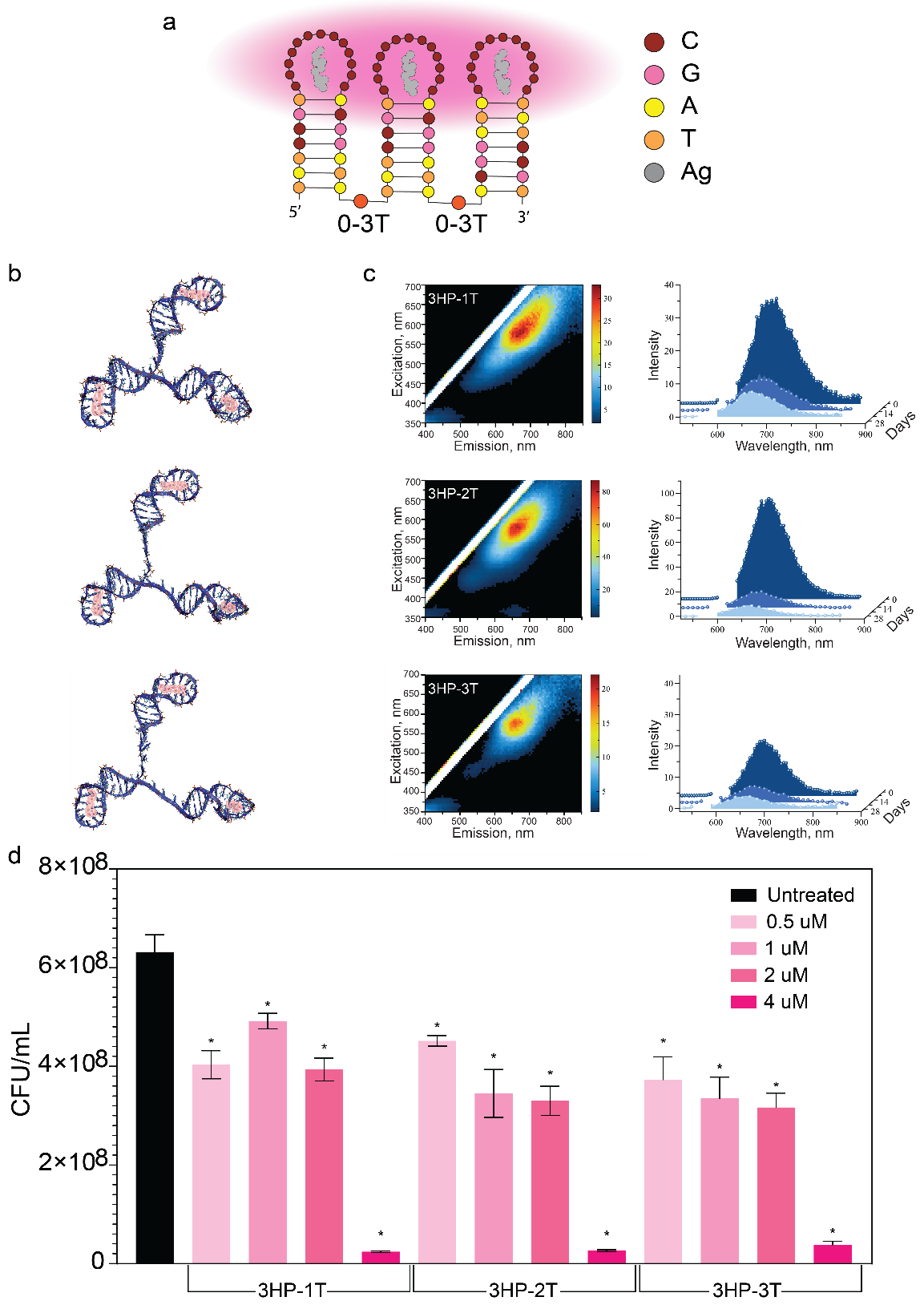

**Supporting Figure S4.** Evaluation of increased flexibility of 3HP DNA-AgNCs with additional thymines. a. Depiction of the 3HP structure and sequence. b. Computational modeling of the structure of 3HP with 1-3 additional thymines. c. Initial 3D excitation-emission spectra with the change in intensity over 4 weeks. d. The anti-bacterial efficacy of DNA-AgNCs at 0.5 to 4 μM.

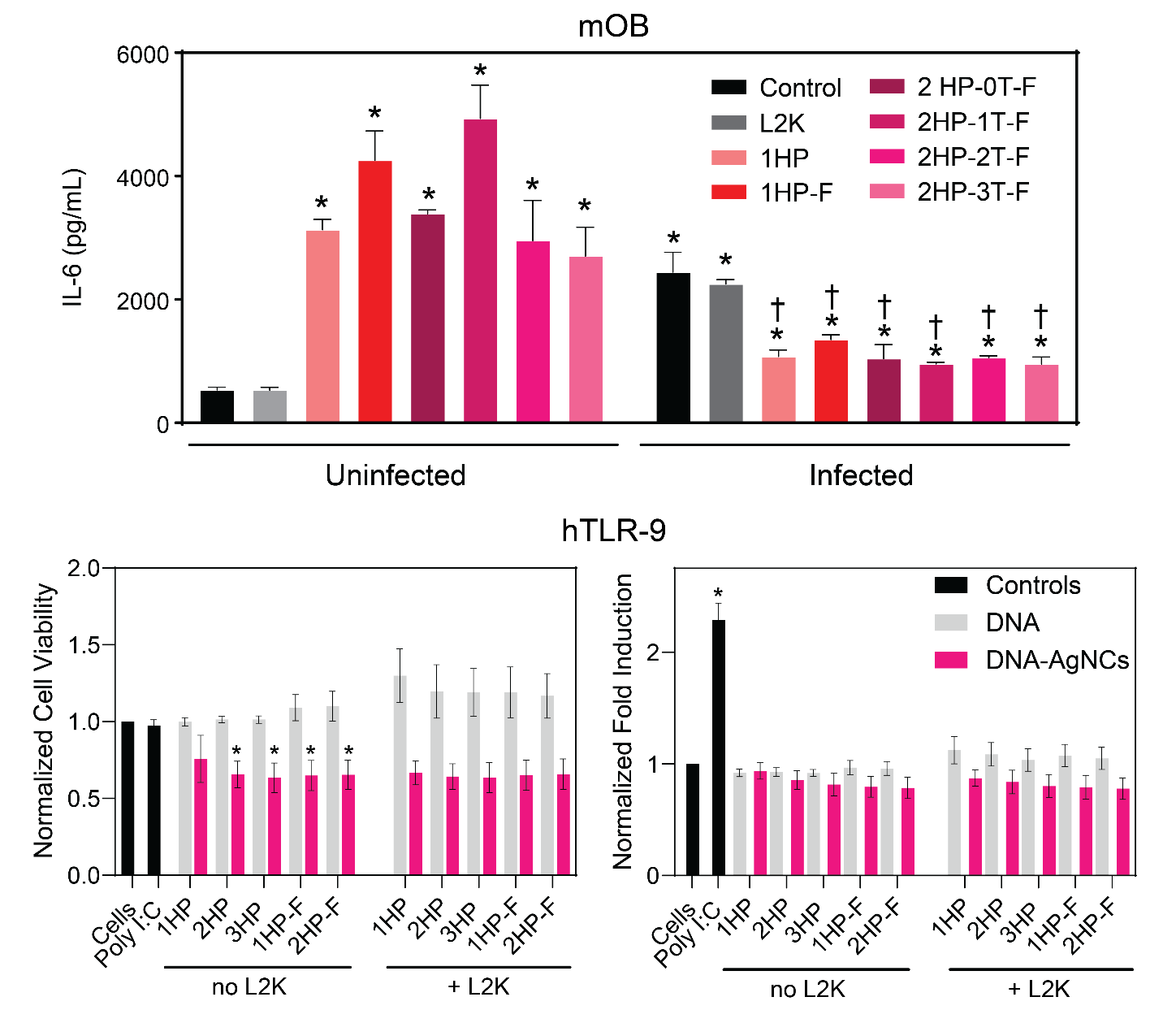

**Supporting Figure S5.** Top: Immunostimulation of primary murine osteoblasts following treatment with with DNA-AgNCs at 0.5 μM in the absence and presence of *S. aureus* infection, 6 hours post-treatment. Asterisks denote significance from the treatment to cells alone. Daggers denote significance between the sample of interest with and without infection. Bottom: Mammalian cell toxicity of multiple hairpin DNA-AgNCs and the immunostimulatory properties of multiple hairpin structures against Human TLR9 Reporter HEK293 cells (hTLR9) at 4 µM DNA when treated with and without Lipofectamine 2000 to evaluate the stimulation of the NF-κB pathway 24 hours post-treatment. Asterisks denote significance from the treatment to cells alone.

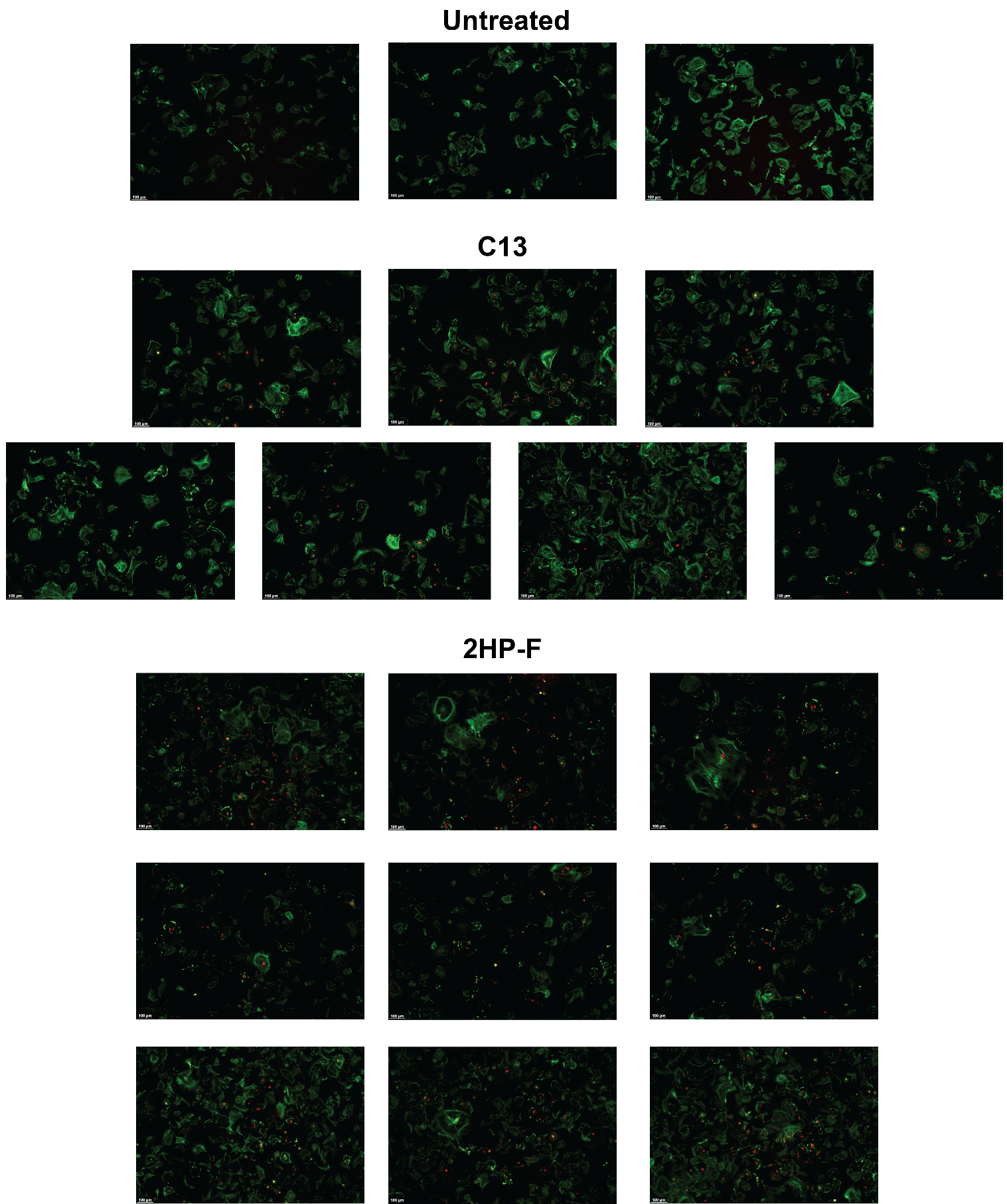

**Supporting Figure S6.** Uptake images of DNA-AgNCs when transfected into primary murine osteoblasts. From top to bottom: osteoblasts without transfected DNA-AgNCs, osteoblasts transfected with C13 DNA-AgNCs, and osteoblasts transfected with 2HP-F DNA-AgNCs.
